# Pharmacodynamic Considerations of Collateral Sensitivity in Design of Antibiotic Treatment Regimen

**DOI:** 10.1101/189381

**Authors:** Klas I. Udekwu, Howie Weiss

**Author notes:** **Correspondence to be addressed to:** Dr. Klas I. Udekwu, Department of Molecular Biosciences, Building F, Room 556 Stockholm, Sweden SE 10691. Equal contribution.

## Abstract

Antibiotics have greatly reduced the morbidity and mortality due to infectious diseases. Although antibiotic resistance is not a new problem, its breadth now constitutes a significant threat to human health. One strategy to help combat resistance is to find novel ways to use existing drugs, even those that display high rates of resistance. For the pathogens *Escherichia coli* and *Pseudomonas aeruginosa*, pairs of antibiotics have been identified for which evolution of resistance to drug A increases sensitivity to drug B and vice versa. These research groups have proposed cycling such pairs to treat infections, similar treatment strategies are being investigated for various cancer forms as well.

While an exciting treatment prospect, no cycling experiments have yet been performed with consideration of pharmacokinetics (PK) and pharmacodynamics (PD). To test the plausibility of this scheme and search for ways to optimize it, we create a mathematical model with explicit PK-PD considerations. We study several possible treatment protocols using pairs of such antibiotics, and investigate the speed of ascent of multiply resistant mutants. Our analyses show that for low concentrations of antibiotics, treatment failure will always occur due to the rapid ascent and fixation of resistant mutants. However, at moderate to high concentrations of some types of bacteriostatic antibiotics with multiday cycling, resistance is prevented from developing and treatment succeeds. This calls for guarded optimism of such treatment protocols whose development can be directed by these types of models.

## INTRODUCTION

The impending era of ubiquitous antibiotic resistance is engendering an urgent search for new therapeutics as well as antibiotic stewardship strategies. Clinical strains have recently been isolated that are completely pan resistant - resistant to all available antibiotics. In lieu of new drugs or vaccines, optimizing therapeutic regimen is critical to continued treatment success. Stewardship approaches incorporating such considerations would ideally reduce the risk of within-host resistance development thus minimizing endemic levels of such strains in the community.

The concept of collateral sensitivity (CS) was described already in the 1950s, when Bryson et al. observed that an *Escherichia coli* strain became hypersensitive to polymyxin B, upon acquiring chloramphenicol resistance. ^1^ They speculated that this collateral sensitivity could be exploited clinically. Recently, research teams have identified drug pairs that exhibit mutual collateral sensitivity (MCS) effects; evolution of resistance to drug A increases sensitivity to drug B and vice versa. MCS entails a reciprocal positive interaction between the two drugs. For the pathogens *Escherichia coli* and *Pseudomonas aeruginosa*, a number of pairs of antibiotics have been identified which exhibit MCS to varying degrees^2-5^ Based on these studies, several groups have proposed antibiotic cycling of two antibiotics (e.g., A -> B -> A -> B -> …, and derivatives thereof) to exploit MCS and to treat infections. ^2^ However, all, except one study were limited to a single exposure to each antibiotic. While recent work has provided some understanding and potential mechanistic causes of such evolved sensitivity, it is unclear if CS is a universally applicable treatment consideration.

For *P. aeruginosa* infections in prophylactically exposed cystic fibrosis, CF, patients, i) gentamicin resistant strains are seen to become sensitive to penicillin due to mutation in a two component system (*pmrB*), and, ii) *nalC* and *mexZ* mutations can confer aminoglycoside sensitivity in beta-lactam adapted strains. ^4^ Data emerging from whole genome sequencing of such isolates points to the need for a thorough exploration of the pharmacodynamic features of such treatment regimen where the CS strategy is employed. Roemhild *et al* cycled two antibiotics at a time in a morbidostat, an in vitro continuous-flow model. ^6^ The morbidostat maintains a population density below a desired threshold by administration of bolus doses of antibiotic at specified optical densities. While a clever and useful device amenable to control and automation, for the potential clinical application, it trivializes the pharmacodynamics and greatly simplifies the underlying modeling. In this case, their model shows rapid ascent of double mutants and experimentally, they find this in most cases. In addition to this, Nichol *et al* recently presented a combinatorial model describing the evolutionary limitations of CS strategy focusing on drugs with the same molecular target; cell wall synthesis. ^7^ They concluded that ‘*collateral sensitivity is contingent on the repeatability of evolution’.*

Theoretical models can guide the testing and implementation of this cycling strategy, in light of the evolutionary trends towards amplification of resistant clones during sequential monotherapies. Using simulated treatment protocols, the plausibility of CS-derived treatment regimen can be evaluated *in silico* and optimized *in vivo*. In this way, the implications for resistance development and the endemic fixation of these alleles can be assessed in ‘real time’ before in vivo testing.

In this manuscript, we analyze the population dynamics of antibiotic cycling with the goal of clinical utility. We utilize clinically relevant parameters for presenting infections; high bacteria densities and antibiotic dosages in line with current treatment protocols. ^8^ Such models combined with PK / PD experiments would guide the clinical applicability of this proposed new treatment paradigm. Our model explicitly incorporates PD / PK parameters for individual strains, their mutation rates and bacterial fitness effects. Implicit within our model is immune control of bacterial density. We examine the sequential application of two drugs, A → B, each exhibiting different PD characteristics, to a hypothetical infection and estimate the rates of resistance ascent or population decline for single and multiply resistant clones in the simulated patient. The simulation results provide a foundation for validating this treatment framework. Time to clearance (of a hypothetical bacterial infection) represents the positive end of the scale, and time to fixation, T_fix_, the negative end which the study focuses on for convenience. The presented framework is well-grounded in the underlying ecological, evolutionary, and pharmacological theory driving collateral sensitivity, and allows us to study the efficacy of treatment achievable under such regimen.

Current examples and proposals of MCS are for two drug cycling and we formulated our original model to study this. It is not only applicable to antimicrobial chemotherapy but also to cancer chemotherapy where investigators have been studying treatment strategies of various cancers based on MCS considerations, although with mixed experimental results.^9^

## MATERIAL AND METHODS

We construct a compartment model where a susceptible population of bacteria, S, is exposed successively to two antimicrobial drugs, A1 and A2, and simulate treatment over a period of 10 days in most cases, and longer where necessary. All strains compete for a limited nutrient and grow at their respective maximum rates in the absence of antibiotic pressure. Resistance confers a fitness disadvantage of 10% for singly resistant mutants with a multiplicative factor of 10% per additional resistance mutation(s). The pharmacokinetics are provided by equations relevant to an *in vitro* kinetic model or chemostat ^10^ at a rate mimicking human glomerular filtration (GFR). The at times neglected pharmacodynamics is incorporated by modulating minimum growth rates relative to each strain’s respective MIC, as opposed to by the MIC alone. Collateral sensitivity is accounted for by switching the MIC of the relevant (pre-exposed) strain at periods corresponding to when the antibiotic is cycled and CS is assumed to be instantaneous. A flow diagram representing the model is presented in Figure 1.

**Figure 1:**
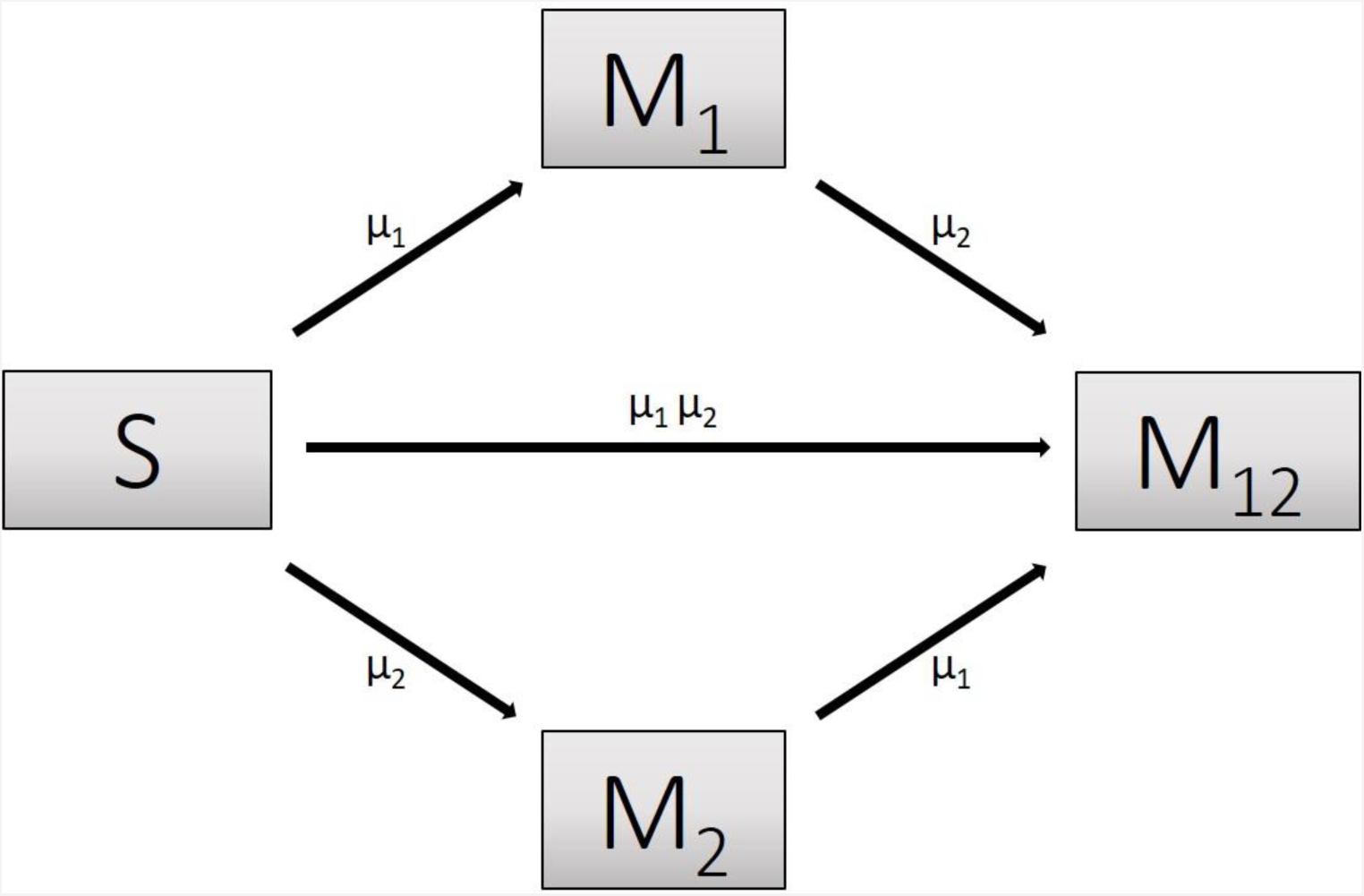
Flow diagram describing the population dynamics of sensitive (S) bacteria exposed to antibiotics and cells resulting from mutation / selection of resistance against antibiotic 1 (M_1_), antibiotic 2 (M_2_), or both (M_12_). Rates of mutation are independent of each other, here described by μ_1_ and μ_2_, and derived from Lee *et al* (2012). ^11^ The rate of double resistance arisal is a product of the individual mutation rates μ_1_ and μ_2_.

### Treatment Model Description

Our treatment protocol consists of exposing a sensitive population of N_0_ = 2 x 10^8^ CFU ml^-1^ of bacteria to each antibiotic as described below. We set the flow rate to 0.2 l h^-1^ and assume a basal mutation rate, μ, of 3.9 x 10^−10^ nucleotide^-1^ generation^-1^, a conservative estimate as compared to the estimated range of μ. ^11^A single mutation is assumed to be sufficient to induce clinically significant resistance and that there is an insignificant likelihood of reversion to full sensitivity. ^12^

For sensitive cells, the MIC is set to 1 μg ml^-1^. When applying drug A or drug B, the MIC of the cells resistant to drug B or drug A respectively, is set to 0.5 μg ml^-1^ and the cells are thus more sensitive than the sensitive cell population. The MIC of resistant cells is always set at 16 μg ml^-1^ prior to switching when antibiotic is cycled. We assume a 10% reduction in fitness by imposing a penalty on maximum growth rate. For each resistance ‘acquisition’ an additive fitness penalty is incurred with the triply resistant isolate being least ‘fit’.

Antibiotics are pulsed at concentration, C μg ml^-1^ (3 ≤ c ≤ 10) into the chemostat for one hour, followed by 5 hours of medium alone. This is repeated four times per day and the antibiotic is switched daily (Pharma7d, 6h). The pharmacodynamic responsiveness of the exposed population to antibiotic concentration is reflected in the Hill coefficient, βwhich defines how ‘rapidly’ the maximum efficacy is attained; the sensitivity of the bacterial population at concentrations close to the MIC. We simulate MCS-guided treatment with daily, and 3 day cycling between antibiotics. We follow each population throughout the treatment period and estimate time to clearance or the time to fixation of multiple drug resistance for either 2 or 3 drug rotations. The inhibitory effect of each antibiotic is reflected in the term, *Ψ*_*min*_, which also reflects the maximal kill rate for each antibiotic. Values for *β* and *Ψ*_*min*_ are derived from experimental studies on *Escherichia coli* and *Staphylococcus aureus*. ^13,14^ We model the intrinsic growth of the population using a Monod model with resource conversion efficiency, *e* = 5 x 10^−7^ μg, and a Monod constant, κ of 0.25 μg ml^-1^ (See Levin et. al, 1977 ^15^). Initially, there are <u>no</u> resistant isolates and we assume that the bacteria do not engage in horizontal gene transfer, a critical assumption for a fair assessment of CS.

Shown below is the coupled system of non-linear differential equations which describe the treatment regimen involving two antimicrobials. We carefully simulate solutions using Matlab. The system is stiff for some parameter values, and we utilize a stiff solver in such cases.

### Mathematical Modeling

For all antibiotics,

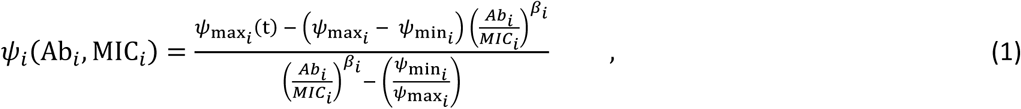

where *i* represents each respective antibiotic (in this iteration, 1 or 2)

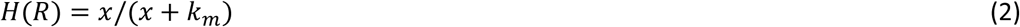

#### Antibiotic 1

*Parameters for dosing Ab*_*1*_

*MIC*_*S*_ = 1; *MIC*_*1*_ = 15; *MIC*_*2*_ = 0.5; *MIC*_12_ = 15

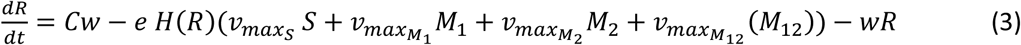

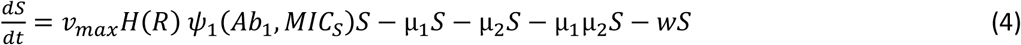

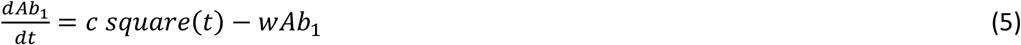

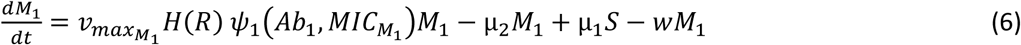

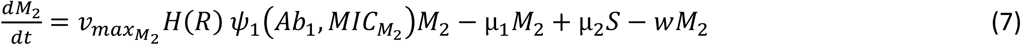

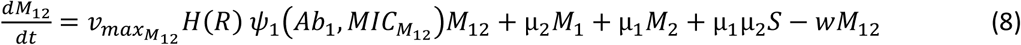

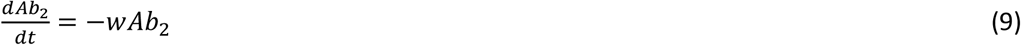

#### Antibiotic 2

*Parameters for dosing Ab*_*2*_

*MIC*_*S*_ = 1; *MIC*_*1*_ = 15; *MIC*_*2*_ = 0.5; *MIC*_12_ = 15

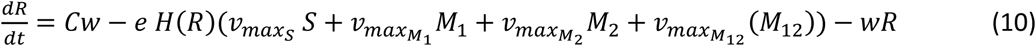

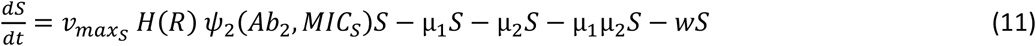

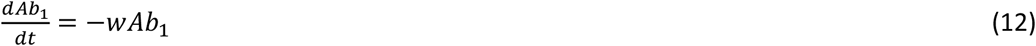

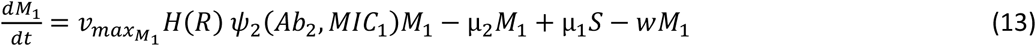

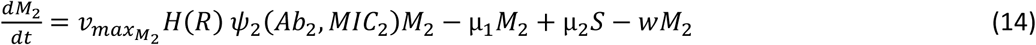

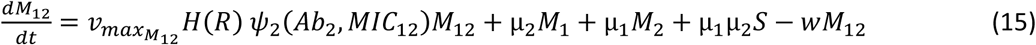

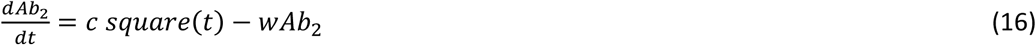

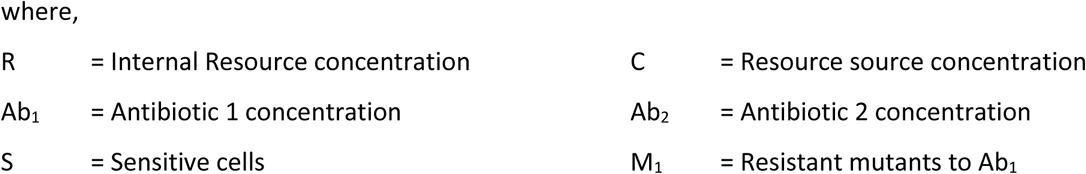

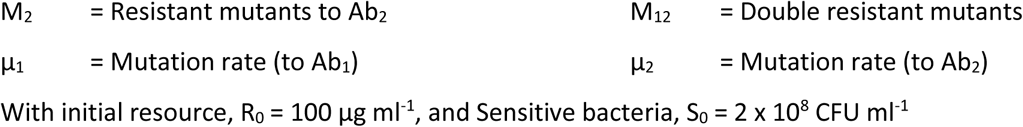

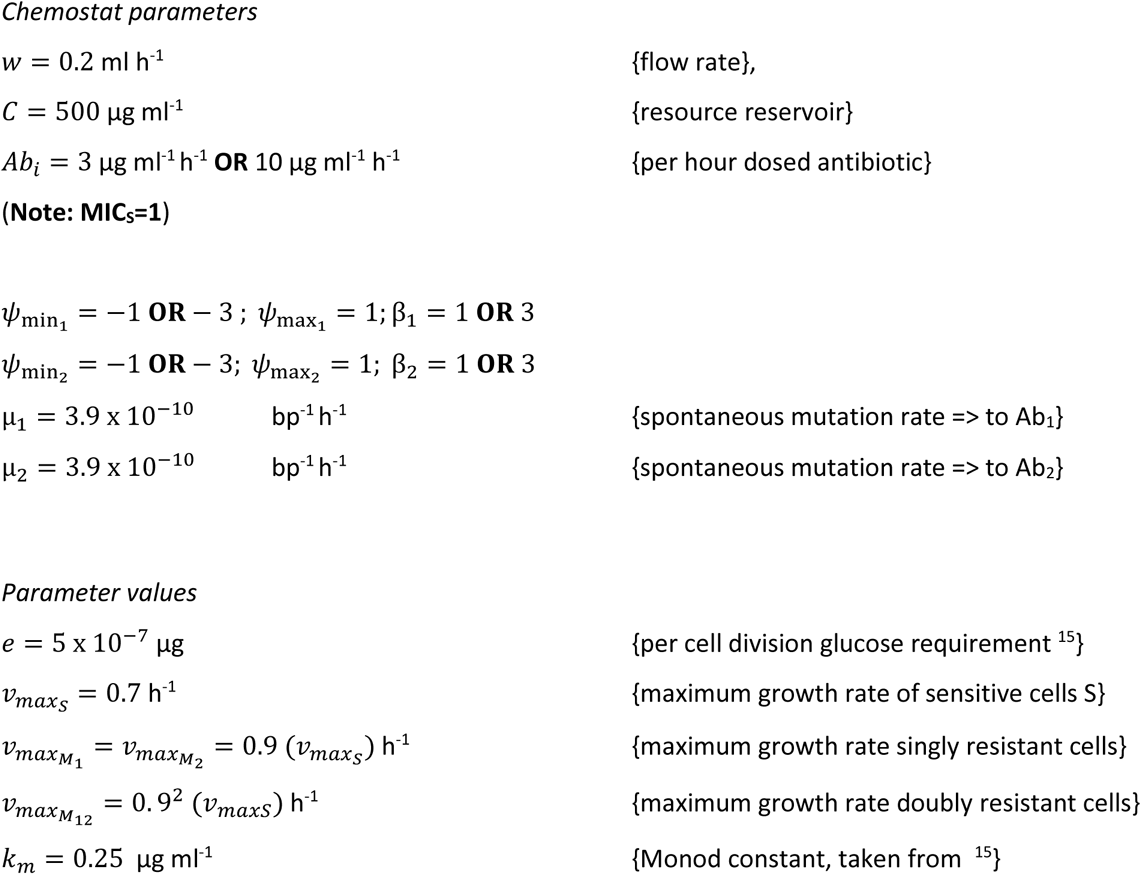

## RESULTS

### In silico assessment of two drug cycling

We simulate treatment with two antimicrobials exhibiting varying PD parameters and encompassing both bacteriostatic (S) and bactericidal (C) drug types (See Table 1 and *Materials & Methods*). We consider both daily cycling and three-day cycling of each antibiotic. We define the time to fixation, T_fix_ as the time between first application of drug 1 and the time at which the double resistant strain exceeds 0.9 N_0_. The simulation results are summarized in Table 1 and displayed in Figure 2.

**Table 1.** Two antibiotic exposure over ten days with daily cycling between Antibiotic 1 and Antibiotic 2 for one day (**ld**) and three day (**3d**) cycling protocols. **C** denotes a bactericidal antibiotic while **S** denotes a bacteriostatic antibiotic. We report resistance fixation T_fix_ in each simulation. For further clarification, see text.

| Min growth rate<br>$\psi_{min_1}$ | Min growth rate<br>$\psi_{min_2}$ | Hill<br>Coefficient $\beta$ | Drug<br>classification | $T_{fix}$ @ $c = 3\mu\text{g ml}^{-1}$<br>(1d, 3d) | $T_{fix}$ @ $c = 10\mu\text{g ml}^{-1}$<br>(1d, 3d) |
| --- | --- | --- | --- | --- | --- |
| -1 | -1 | 3 | S/S | 67h, 102h | 95h, 113h |
| -1 | -3 | 3 | S/C | 67h, 101h | 90h, 107h |
| -3 | -1 | 3 | C/S | 68h, 101h | 90h, 112h |
| -3 | -3 | 3 | C/C | 68h, 101h | 87h, 107h |
| -1 | -1 | 1 | S/S | 96h, 115h | 438h, >1000h |
| -1 | -3 | 1 | S/C | 90h, 110h | 294h, 1198h |
| -3 | -1 | 1 | C/S | 90h, 115h | 233h, 1124h |
| -3 | -3 | 1 | C/C | 85h, 109h | 237h, 523h |

**Figure 2.**
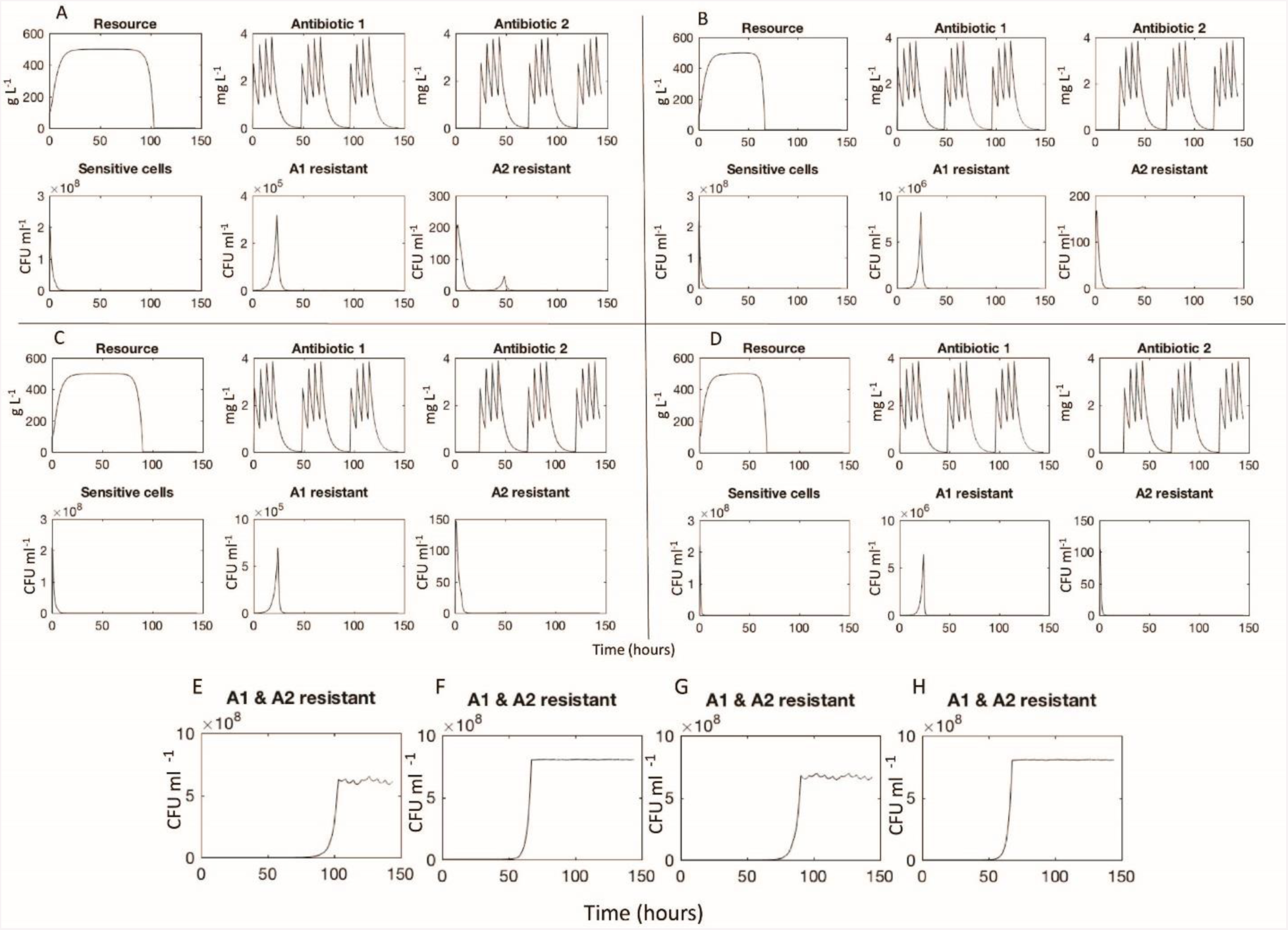
Selected simulations of our mathematical model showing changes in resource and antibiotics concentrations as well as population densities or sensitive and mono-resistant cells. Each panel A-D represents different runs for input antibiotic concentrations of 3 μg ml^-1^ (**A**: *β* = 1, S / S; **B**: *β* = 3, S / S; **C**: *β* = 1, C / C; **D**: *β* = 3, C / C). **E-H** represent the population densities of doubly resistant cells => T_fix_ for runs **A-D** respectively.

For low antibiotic dosing (∼ 3x MIC of sensitive cells), fixation of double mutants, i.e. resistance to both applied antibiotics occurs rapidly, T_fix_ ranging between 67 hours and 115 hours. This occurred despite the increased susceptibility imposed by CS theory. The fixation of doubly resistant strains is seen to be invariable, although it takes 50% longer when drugs are cycled every three days than when cycling daily. The use of drugs with smaller Hill coefficients (β) leads to a 25% delay in fixation. Less significant are the differences between the various combinations of static and cidal drugs. For drugs with small Hill coefficients however, there is a 5% delay in fixation when one static drug is used and a further 5% delay when both cycled drugs are bacteriostatic.

For moderate to high dosing schemes (∼ 10x MIC of sensitive cells), for drugs having large β, doubly resistant mutants fixate rapidly, within four days of treatment initiation. This is not the case for drugs with small *β* values with which resistance only emerges (and fixates) long after a traditional clinical treatment regimen is completed, i.e. > 7-10 days. Tfix also decreases when using two bactericidal (C / C) drugs as compared to other combinations of cidal and static drugs (C / S or S / C). The best strategy of all from our simulations is the use of two bacteriostatic drugs (S / S) with a low β, while cycling them every three days. Under these conditions, no resistance is observed for any conceivable infection treatment duration, which we interpret as meaning that such regimen with MCS guiding protocol may actually inhibit the development of resistance.

Lastly, we note that significant short duration population spikes (> 10^5^ CFU ml^-1^) occurred for some singly resistant strains.

## DISCUSSION

The emerging crisis that is pan antibiotic resistance, compounded by a dearth of new antibiotics in the development pipeline ^16^ is necessitating new approaches to chemotherapy to minimize resistance emergence and dissemination. Many clinical treatment strategies have been proposed, ranging from combinatorial therapy to antibiotic cycling incorporation in the treatment of everyday infections in order to mitigate resistance emergence. Antibiotic cycling can be implemented either on the level of individual patients or the institutional unit. Earlier modeling studies of unit cycling cast doubts on the utility of this strategy ^17^ and results of hospital trials have been mixed.

The MCS approach is an exciting treatment option but multiple cycle testing with clinically relevant PK/PD consideration has not been explored. Our mathematical model allows us to do precisely this, focusing on the PD properties relevant to treatment of a hypothetical infection. Our modeling strategy not only considers the classical <u>M</u>inimum <u>I</u>nhibitory <u>C</u>oncentration but also the dose responsiveness of a population of sensitive and resistant bacteria in a resource-limited setting. This provides a broader perspective than the usual one parameter, MIC, summary on the PD.

Several drug pair combinations have been identified as exhibiting MCS as earlier stated. Results from WGS based studies have identified two possible mechanisms steering MCS: i) intragenic suppression of resistance, ^7^ and, ii) epistasis derived reversal of resistance (towards the other drug). ^2^ These are based on both the mutation target size and the breadth of the path each adaptive walk has traversed. Intragenic suppression is limited by the smaller number of mutations that can lead to an MCS phenotype for a related antibiotic while epistatic suppression is more dependent on an underlying genetic diversity. These are also discussed in ^7^ and the observed saddlepoint in their published evolutionary landscape points to potential treatment failure; evolutionary unpredictability as such is not considered herein.These only serve to bolster the importance of our comprehensive pharmacodynamic study.

Pharmacodynamic properties of antibiotics are broadly delineated as bacteriostatic or bactericidal for ease of pharmacological understanding. Most bacteriostatic drugs are weakly -cidal and while bactericidal drugs are often preferred to bacteriostatic drugs, in immunocompetent patients, there is no advantage in using the former. From our simulations, it appears that when exploiting MCS, the bacteriostatic drugs are superior to their -cidal counterparts, as they inhibit resistance development. To our knowledge, this is the first report or observation of such, emerging albeit from an *in silico* study. As presented in the Results section, we observe the rapid fixation of resistance in all cases of low drug concentration exposure, underlining the importance of adequate dosage. The large population sizes commonly associated with infections, also considered in our study, would require an immunocompetent host (patient) to clear. At first glance, our model lacks both innate and adaptive immunity (approached in ^18^) but the innate immunity is implicit in form of resource concentration limitation on population size.

From the estimates of T_fix_ the single factor most significant in the early fixation of multiple resistance using this MCS protocol is the cycling period (*cv* 67 h vs. 110 h for one day cycling and three day cycling respectively). Following this is the effect of the Hill coefficient, β, on T_fix_ which, in line with, ^19^ challenges the idea that increased clearance rates minimize the selection and fixation of resistant isolates. The increased selective pressure imposed by higher order bactericidal drugs leads to a more rapid fixation of resistance in the population. Our model explains this from a nutrient competition perspective; as antibiotic 2 is being applied, the killing of the sensitive cells and those resistant to antibiotic 1, results effectively in reducing the total population density and thus competition is relaxed.

This is also true for the other antibiotics and evolution of doubly resistant mutants is facilitated by the large population sizes being acted upon by the imposed (conservative) mutation rate in our model (See Figure 2). Once doubly resistant, this collateral sensitivity is lost and overall fitness of the population is maximized under (either Ab_1_ or Ab_2_) antibiotic pressure.

It is known that the efficacy of beta lactam antibiotics is highest at concentrations around five times the MIC and the results obtained from our low dose simulations are thus interesting for this highly clinically utilized class of antibiotics. The PD parameter *β* describes the sensitivity of the antibiotic to changes proximal to the point of inflection (zero growth) in the graph of the Hill function (See model description). This plays a vital role in the success of MCS-guided treatment regimen although it is not evident *á priori* (See Figure 3). As is seen upon replotting the data to reflect the effect of *β* on our estimate of efficacy, the fixation time for bactericidal drugs in particular is heavily dependent on the drug dosage especially for the 3d cycling protocol. These are strong indicators of the importance of drug choice under such treatment regimen consideration.

**Figure 3:**
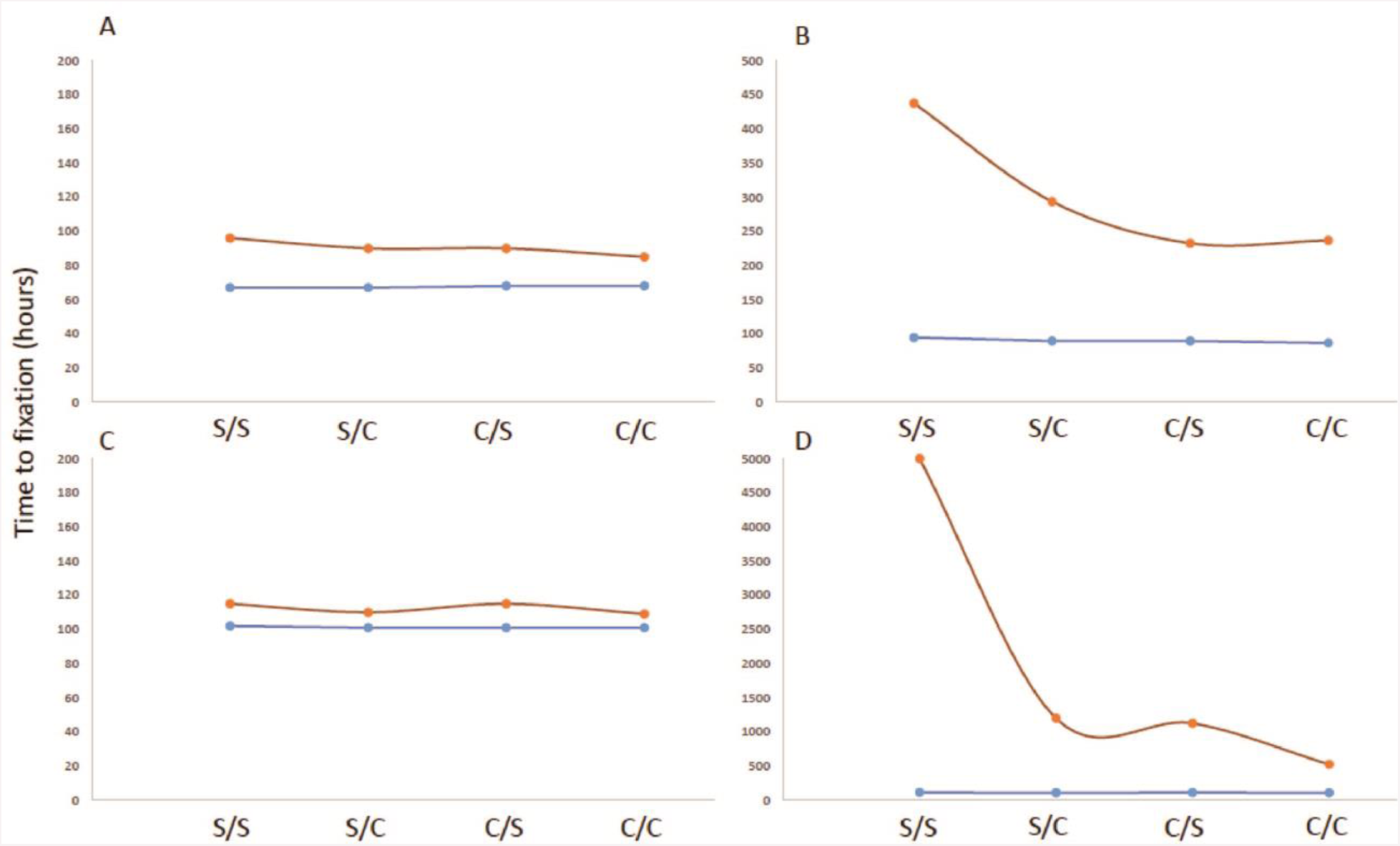
Categorical plot showing the effect of parameter values *β* and Ψ^*min*^ on fixation time, *T*^fix^, for two drug concentrations; 3μg ml^-1^ (Panels ***A*** & ***C***) or 10μg ml^-1^ (Panels ***B*** & ***D***) of combinations of cidal (*C*) and static (*S*) drugs (*x-axes*) were simulated with *β* = 1 (*Red*) or *β* = 3 (*Blue*) and cycled either every day (Panels ***A*** & ***B***) or every 3 days (Panels ***C*** & ***D***).

In vivo, the immune system works hand in hand with antibiotics to clear the infection. In some of our simulations, the single and double mutants remain at low enough densities to be easily cleared by circulating immune cells. However, when fixation of double mutants occurs within a few days, the likelihood of treatment failure would by definition, increase. Even in cases where fixation does occur, the immune system, unaffected by AMR state may still clear. As such our model only serves as a cautionary tale; heterogeneity in immunity and an immunocompromised state could exacerbate the likelihood of treatment failure, as would antimicrobial heteroresistance, inoculum effects and high frequency of AMR bacteria in the original infection.

In addition, horizontal gene transfer confounds the model’s predictability as the rate of plasmid-mediated resistance transmission, especially conjugable plasmids, is considerably higher than the replication dependent vertical propagation of point mutations. We do not consider plasmid-based transmission of antimicrobial resistance genes and neither do we include any resistant mutants initially, only considering de novo mutations conferring resistance. These mutations we assume occur independently of one another. Under continuous antibiotic exposure, selection drives the system towards a fitness maximum through resistance compensation which is favored over reversion ^12^.

The modeling framework we present in this manuscript treats two drug cycling but has already been extended to three drug cycling in anticipation of triplet combinations that may be identified in due course. Before clinical deployment however, the utility of MCS must be explored experimentally, rationalized physiologically, and further tested for generality. Mathematical models of this sort will play a crucial role in establishing the applicability and suggesting modifications to this and other proposed treatment regimen.

## ACKNOWLEDGEMENTS

The authors acknowledge the KITP program on Ecology and Evolutionary Biology 2017 (NSF Grant No. PHY17-48958, NIH Grant No. R25GM067110, and the Gordon and Betty Moore Foundation Grant No. 2919.01)

